# Glutton: large-scale integration of non-model organism transcriptome data for comparative analysis

**DOI:** 10.1101/077511

**Authors:** Alan Medlar, Laura Laakso, Andreia Miraldo, Ari Löytynoja

**Affiliations:** Institute of Biotechnology, University of Helsinki, Finland; Center for Macroecology and Evolution, University of Copenhagen, Denmark

## Abstract

High-throughput RNA-seq data has become ubiquitous in the study of non-model organisms, but its use in comparative analysis remains a challenge. Without a reference genome for mapping, sequence data has to be *de novo* assembled, producing large numbers of short, highly redundant contigs. Preparing these assemblies for comparative analyses requires the removal of redundant isoforms, assignment of orthologs and converting fragmented transcripts into gene alignments. In this article we present Glutton, a novel tool to process transcriptome assemblies for downstream evolutionary analyses. Glutton takes as input a set of fragmented, possibly erroneous transcriptome assemblies. Utilising phylogeny-aware alignment and reference data from a closely related species, it reconstructs one transcript per gene, finds orthologous sequences and produces accurate multiple alignments of coding sequences. We present a comprehensive analysis of Glutton’s performance across a wide range of divergence times between study and reference species. We demonstrate the impact choice of assembler has on both the number of alignments and the correctness of ortholog assignment and show substantial improvements over heuristic methods, without sacrificing correctness. Finally, using inference of Darwinian selection as an example of downstream analysis, we show that Glutton-processed RNA-seq data give results comparable to those obtained from full length gene sequences even with distantly related reference species. Glutton is available from http://wasabiapp.org/software/glutton/ and is licensed under the GPLv3.

## 1 Introduction

High-throughput RNA sequencing (RNA-seq) is used to efficiently characterise transcriptome structure, quantify gene expression and infer post-transcriptional modifications such as alternative splicing (Wang *et al.*, 2009). As RNA-seq captures gene regions, it has become popular for quickly and cheaply producing data for evolutionary studies. Indeed, large-scale efforts such as the “1,000 plants project” (Matasci *et al.*, 2014) and the “1K insect transcriptome evolution” (Misof *et al.*, 2014) projects have opted for transcriptomes, instead of whole genome sequences, to comprehensively sample their respective clades for comparative analyses. While this versatility has made RNA-seq ubiquitous in the study of non-model organisms, its application remains challenging. For example, non-model organisms lack reference genomes, necessitating *de novo* assembly. Lowly expressed transcripts can be fragmented or otherwise misassembled due to sequencing errors and even perfectly assembled transcripts lack orientation information and include untranslated regions (UTRs).

In evolutionary studies, transcripts must be aligned such that the inferred character homologies reflect the evolutionary history of each gene. Erroneous sequence alignments can lead to incorrect inferences with, for example, branch-site (Fletcher and Yang, 2010) and site-wise (Jordan and Goldman, 2012) tests for positive selection. Several studies have demonstrated that the evolutionary alignments produced by PRANK (Lo¨ytynoja and Goldman, 2008) more accurately reflect evolutionary history compared to other methods and are therefore more appropriate for comparative studies (Fletcher and Yang, 2010; Privman *et al.*, 2012; Jordan and Goldman, 2012). PRANK implements the phylogeny-aware alignment algorithm (Lo¨ytynoja and Goldman, 2005) which uses phylogenetic information to model substitution probabilities and distinguish between insertions and deletions. More recently, the concept of evolutionary alignment has been extended to the alignment of partial-order sequence graphs (Lo¨ytynoja *et al.*, 2012). Using this approach, PAGAN can align fragmented sequences by extending an existing alignment, which has already been employed in diverse contexts (e.g. Medlar *et al.* (2014); Veidenberg *et al.* (2015)). In the analysis of RNA-seq data, alignment extension enables reference data from related species to be used as a backbone to process the assemblies, providing locality, orientation and homology information. However, a complete RNA-seq data set still requires extensive processing including assigning contigs to reference genes, support for parallelism and suitable post-processing such as the removal of UTRs.

Here we describe Glutton, a new transcriptome scaffolding approach for RNA-seq data from non-model organisms that is intended for use together with downstream comparative analysis methods. Glutton leverages reference data from EnsemblCompara GeneTrees (Vilella *et al.*, 2009), taking advantage of evolutionary information to distinguish between paralogous genes. It accurately scaffolds contigs where there is either insufficient overlap or coverage for *de novo* assembly to connect discontiguous sequences. While transcriptome assemblers can produce very many transcript isoforms for a given gene, Glutton aims to generate a single consensus sequence per gene for each species under investigation by integrating sequencing depth information to merge transcript isoforms.

## 2 Results

Glutton is comprised of three distinct stages: build, align and scaffold (see Figure 1). The most common usage requires only the last two stages, as reference databases can be pre-built.

**Figure 1:**
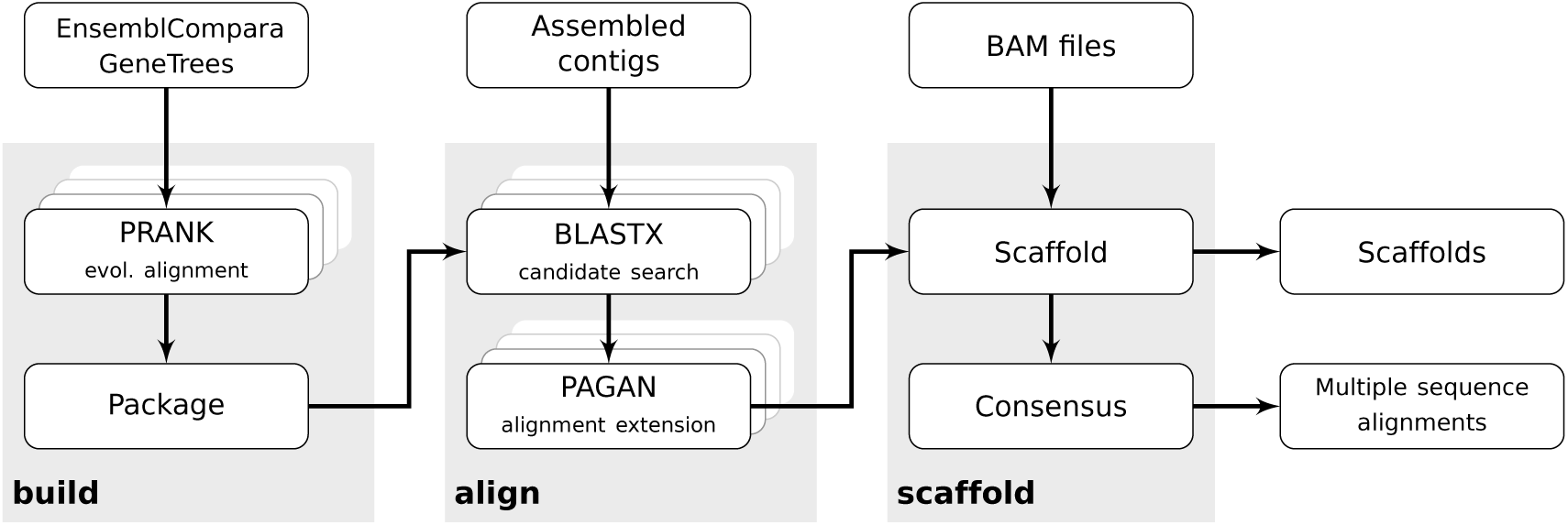
Glutton has three stages: build, to create a reference database from EnsemblCompara GeneTrees; align, to assign contigs to reference genes and perform alignment extension per gene family; and scaffold, to generate scaffolds and consensus alignments for downstream analysis. Overlapping boxes indicate use of multiple CPUs.

**Build:** Glutton requires a reference database that contains evolutionary alignments of all CDS sequences from a chosen reference species. In the current implementation, we use homology information from EnsemblCompara GeneTrees to group reference sequences together into gene families (which, as we use only one reference species, are sets of paralogs). PRANK is used to perform a translated alignment of each gene family. The resulting sequence alignments and phylogenetic trees are packaged into a single database file.

**Align:** Glutton’s approach is based on phylogeny-aware alignments provided by PAGAN. As input, Glutton accepts an arbitrary number of transcriptome assemblies from multiple species, possibly including multiple samples per species. In the alignment step, Glutton bins the input contigs together with putative homologous candidate genes from a given reference database. For each input contig we use BLASTX (Camacho *et al.*, 2009) to perform a translated local alignment and accept the top BLAST hit as the candidate gene for that contig. As candidate genes are obtained using local alignment, paralogs of that candidate gene might be better matches when using global alignment. To allow for this uncertainty, instead of aligning contigs directly with candidate genes, we use PAGAN to extend the full reference alignment of each gene family with the contigs assigned to its members.

**Scaffold:** In the scaffolding step, Glutton post-processes the sequence alignments generated by PAGAN. Contigs within the alignments are filtered and those with insufficient identity and overlap with the reference transcripts are discarded. The remaining contigs are merged into scaffolds if there is sufficient evidence from the reference gene and no contradictory information, such as gaps in the alignment between two contigs. Two contigs are merged if, for example, the overlap between the contigs is lower than the *k*-mer length used during assembly or two disconnected contigs are both highly similar to the reference gene. In contrast, contigs identified by the assembler as different isoforms of the same gene should not be merged into a scaffold. Glutton currently supports alternative isoforms inferred by Oases (Schulz *et al.*, 2012), SOAPdenovo-Trans (Xie *et al.*, 2014), and Trinity (Grabherr *et al.*, 2011).

Along with scaffold sequences for each sample, Glutton produces multiple sequence alignments of homologous sequences across different samples. In this, the objective is to maximize the number of truly homologous alignment positions, possibly accepting that the sequences in the resulting alignment are composites of multiple different transcripts. Consensus sequences are produced using sequencing depth information from mapping raw reads to contigs. For each gene, contigs are ordered by average depth and merged sequentially from lowest to highest, allowing contigs with greater depth to overwrite those with lower. The resulting sequences are optionally trimmed by cutting their 5’ UTR sequences at the start codon and truncating the 3’ ends at any stop codons. As downstream analysis methods expect complete genes, Glutton discards alignments with low sequence coverage. By default, only alignments with more than 90% of the reference gene covered are retained.

Glutton’s ability to produce high quality alignments is contingent on the availability of an appropriate reference species and is affected by other factors such as the transcriptome assembler used.

### 2.1 Effect of reference divergence

To understand the effect of divergence time between study and reference species, we selected three study species from Ensembl Metazoa (Kersey *et al.*, 2016): *Acyrthosiphon pisum*, *Anopheles darlingi* and *Drosophila melanogaster*. *A. pisum* was chosen because it is the species with the highest number of genes (36,195), *A. darlingi* because it has the least (10,457) and *D. melanogaster* because it has a middling number of genes (13,918) and the highest number of closely related species (11 from the genus *Drosophila*). For reference species, we used 55 species from Ensembl Metazoa (release 25). As the combination of data set and transcriptome assembler are potentially confounding factors, instead of assembled contigs we used the complete reference transcriptomes for the three study species as input to Glutton. We used EnsemblCompara to assess orthology. Divergence times were taken from the TimeTree database (Hedges *et al.*, 2015). At the time of writing, divergence times were not available for *Megaselia scalaris*, *Trichinella spiralis* and *Trichoplax adhaerens*, so these were omitted from figures. In all figures, divergence times have been jittered *±*20 million years (mya) to improve legibility.

Results for the 165 combinations (3 x 55) of study and reference species show that the number of alignments output is positively correlated with the number of genes in the study species (Figure 2). The total number of alignments decays exponentially with divergence time between study and reference species. The proportion of genes recovered, however, is higher for study species with fewer genes across the complete range of divergence times (Figure S2).

**Figure 2:**
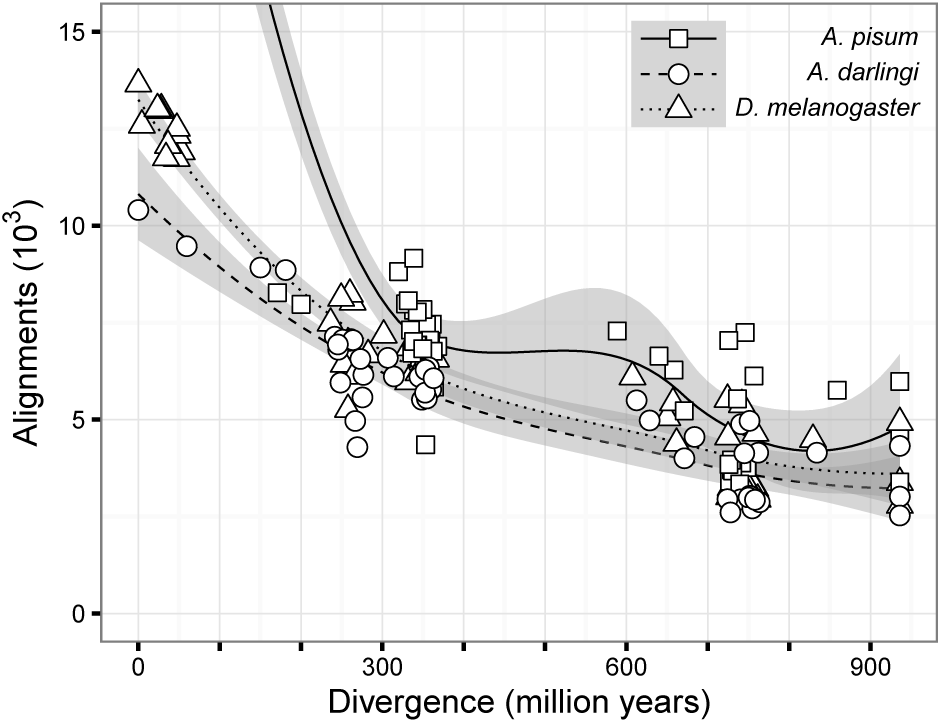
Number of alignments output as a function of divergence time. The three study species have different numbers of genes: 36,195 in *A. pisum*, 10,457 in *A. darlingi* and 13,918 in *D. melanogaster*. For clarity *A. pisum* using itself as reference has been omitted. The complete graph is shown in Figure S1.

Ortholog selection is highly sensitive in closely related species (divergence time *<* 50 mya) with sensitivity scores from 0.95 (Figure 3). In contrast, for distantly related species, the specificity of ortholog selection is high. In the case of *A. pisum*, specificity is consistently greater than 0.9 with all reference species (Figure S3). The proportion of error-free alignments decays with divergence time at a faster rate than sensitivity, reaching a plateau after 350 mya (Figure 4). An alignment is considered “error-free” if all of the constituent query sequences are orthologous to the reference gene according to EnsemblCompara (see Figure S4 for ortholog selection accuracy) and there are no many-to-one ortholog relationships resulting in chimeras. This definition sidesteps whether the alignment itself contains false homologies, despite correct ortholog assignment. The height of the plateau is, like sensitivity, negatively correlated with the number of genes in the study species. In the case of *A. pisum*, the high number of genes results in many-to-one orthology relationships, which in turn cause Glutton to generate erroneous alignments. For *A. darlingi*, the species with the fewest genes tested, the proportion of error-free alignments does not dip below 0.7, even with reference species separated by 900 mya.

**Figure 3:**
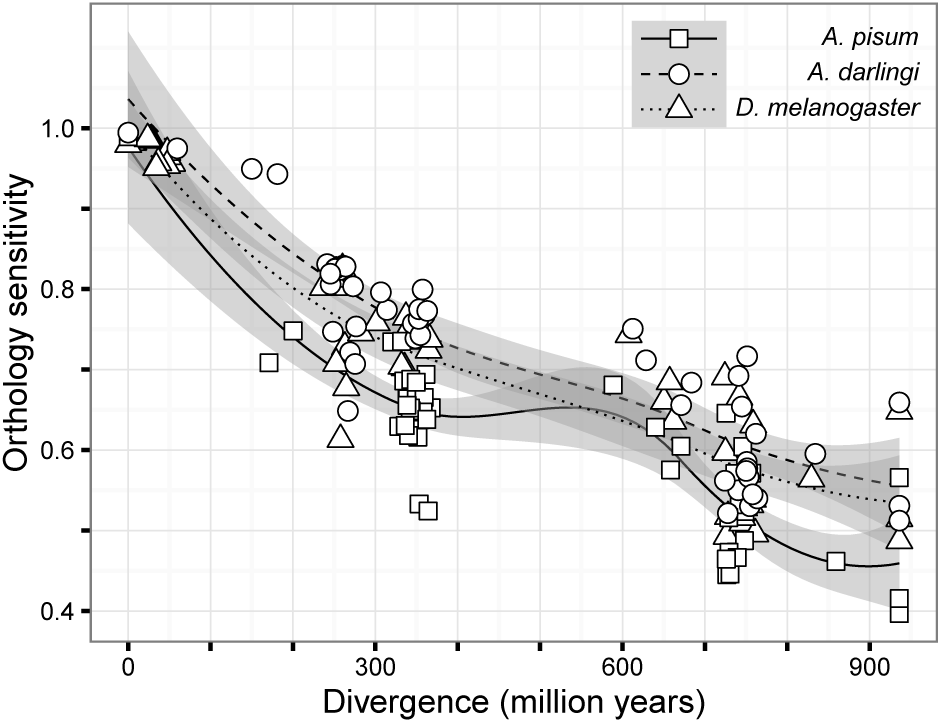
Sensitivity of ortholog selection as a function of divergence time.

**Figure 4:**
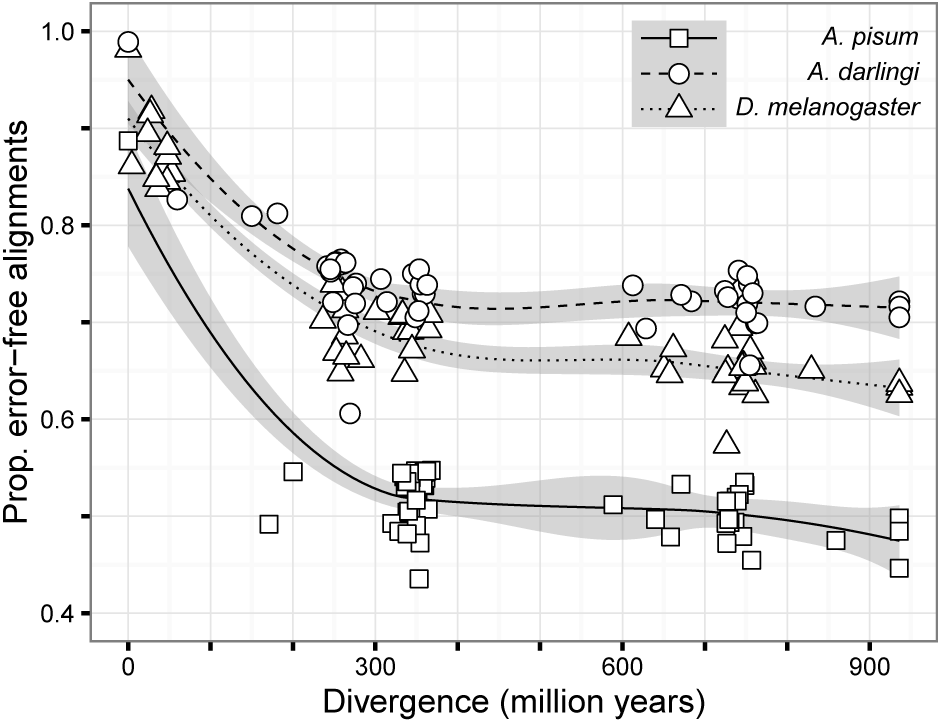
Proportion of non-chimeric genes with the correct orthology as a function of divergence time.

### 2.2 Effect of assembler

Widely used transcriptome assemblers are based on de Bruijn subgraphs and use paired-end reads spanning splice site junctions to infer alternative splicing isoforms. Despite similarities in the underlying methods, studies have observed large differences in, for example, isoform detection sensitivity (Schulz *et al.*, 2012). As Glutton relies on reference data consisting of canonical transcripts and not complete gene models (complete gene models are rarely available for non-model species), we sought to understand how different transcriptome assemblers affect performance. We tested four widely used transcriptome assemblers: Oases (Schulz *et al.*, 2012), SOAPdenovo-Trans (Xie *et al.*, 2014), TransABySS (Robertson *et al.*, 2010) and Trinity (Grabherr *et al.*, 2011). We retrieved RNA-seq data from four *D. melanogaster* samples (accessions SRR166807, SRR166808, SRR166809 and SRR166810) from the Sequence Read Archive (SRA) (Kodama *et al.*, 2012) which were filtered for quality and assembled using the four different assemblers. For each assembly, raw reads were mapped to assemblies using BWA (Li, 2013). The combination of four RNA-seq data sets and four different assemblers gave us sixteen separate samples. We processed all samples with Glutton using 55 reference species from Ensembl Metazoa (release 25) (see Supplemental methods for further details).

From the resulting data, we looked at two metrics to compare the assemblers: the number of alignments output by Glutton and the proportion of those alignments that were error-free (i.e. contigs assigned to correct ortholog and non-chimeric consensus sequence). In the previous experiment we assessed orthology directly using EnsemblCompara. Here we indirectly assessed orthology by assuming that whichever gene a contig was aligned to in *D. melanogaster* was the ground truth and compared the orthology of that *D. melanogaster* gene with the assigned genes from the other references. To be included in this analysis, alignments were required have at least 90% coverage of a reference gene, at least 30% protein identity to the same gene and be at least 100 bp long. Early on, we identified SOAPdenovo-Trans as having poor performance due to its own scaffolding procedure inserting gaps (Ns) that interfered with the Glutton alignment step. We therefore reran SOAPdenovo-Trans in single-end read mode to avoid this problem, but also show the original paired-end results.

Table 1 shows the results from sample SRR166808 with reference species: *Drosophila grimshawi* (divergence from *D. melanogaster*, 47.6 mya), *Zootermopsis nevadensis* (348 mya) and *Mnemiopsis leidyi* (936 mya). The number of contigs produced by different assemblers varies widely, ranging from under 42,000 to nearly 185,000, however, the number of alignments output by Glutton are very similar irrespective of assembler. While there are differences between assemblers, the reference species accounts for a majority of the variance with both the number of alignments and the correct proportion decreasing rapidly with increasing divergence time. We note that the proportion of error-free alignments from RNA-seq data is higher than for reference transcripts (Figure 4). This difference can be attributed to the transcriptome assemblers merging parts of different genes with high similarity, calling them different isoforms of the same gene. Despite the number of alignments varying widely between reference species, gene coverage was more dependent on sequencing depth than any other variable. For example, for the three reference species shown in Table 1, the median gene coverage only varied between 99.1-99.7% (data not shown).

**Table 1:** Transcriptome assembler and Glutton statistics for sample SRR166808 with example close, medium and distantly related reference species (*D. grimshawi*, *Z. nevadensis* and *M. leidyi*). *\** = rerun as single-ended reads.

|  | Contigs | Scaffolds | *D. grimshawi* |  | *Z. nevadensis* |  | *M. leidy* |  |
| --- | --- | --- | --- | --- | --- | --- | --- | --- |
|  |  |  | Alignments | Correct (%) | Align. | Corr. | Align. | Corr. |
| TransABYSS | 184,686 | n/a | 6,718 | 92.1 | 3,668 | 85.2 | 1,447 | 78.9 |
| Oases | 85,843 | 22,544 | 6,604 | 88.7 | 3,579 | 80.3 | 1,370 | 72.0 |
| Trinity | 41,814 | 25,882 | 6,721 | 92.7 | 3,730 | 86.2 | **1,545** | 80.5 |
| SOAPdenovo-Trans | 45,047 | 43,497 | 5,979 | **94.2** | 3,536 | **87.7** | 1,490 | 81.0 |
| SOAPdenovo-Trans* | 53,277 | n/a | **6,878** | 93.3 | **3,797** | 87.3 | 1,543 | **81.7** |

To gain a deeper understanding of how the choice of assembler affects the performance, we modelled the relationship between the number of Glutton alignments and the assembler as a generalised linear model (GLM) controlling for both sample and divergence time. Briefly, we assumed the response variable to be negative binomially distributed and used a log link function. We tested each term in the model for significance using the likelihood ratio test versus reduced models and all terms were significant. *Post hoc* pairwise comparisons between all assemblers showed SOAPdenovo-Trans with paired-end data produced significantly less alignments compared to Trinity and SOAPdenovo-Trans with single-end data (both *p <* 0.001, Tukey’s test). All other pairwise combinations were non-significant, suggesting there is little difference between a majority of the assemblers tested with respect to the number of alignments. To assess the quality of results, we modelled the proportion of error-free alignments as a function of the assembler, again controlling for sample and divergence time. We used beta regression together with the logit transform (Ferrari and Cribari-Neto, 2004), all terms in the model were significant by the likelihood ratio test. *Post hoc* analysis of the assemblers revealed all pairwise comparisons were statistically significant (*p <* 0.01) with the exception of TransABySS/Trinity (*p* = 0.77) and SOAPdenovo-Trans single/paired (*p* = 0.64, Tukey’s test). Assemblies from SOAPdenovo-Trans (both modes) consistently produced the highest proportion of correct alignments, followed by a tie between Trinity and TransABySS. Assemblies from Oases produced the lowest proportion of correct alignments. Overall, the best assembler for use with Glutton is SOAPdenovo-Trans with single-ended data as it produced the highest number of alignments, the highest proportion of which were error-free.

The reference genomes used from Ensembl Metazoa are at varying stages of completeness and use different sequencing technologies, all of which may have an impact on Glutton’s performance. Our modelling can be used to identify outliers by comparing the observed proportion of error-free alignments from our experiments with the predictions made by our beta regression model (in-sample accuracy). We identified *Melitaea cinxia* (Ahola *et al.*, 2014) as a clear outlier that consistently underperformed compared to model predictions (Figure 5). While it is unclear what would cause a single reference to cause Glutton to under perform versus predictions, we note that *M. cinxia*, of the species we tested, is the most recently accepted genome into Ensembl Metazoa and we speculate that the ortholog definitions may be incomplete, raising the error rate.

**Figure 5:**
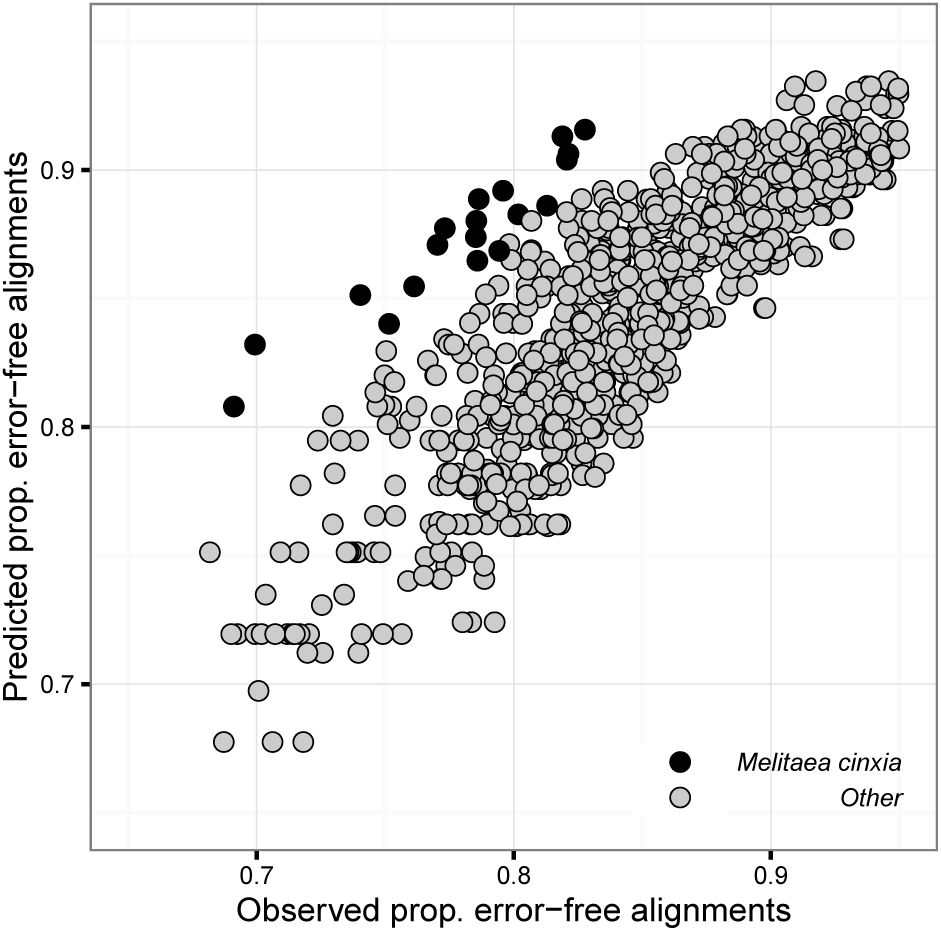
Observed proportion of error-free alignments versus model predictions. *M. cinxia* is highlighted as consistently underperforming compared to model predictions.

### 2.3 Comparison with alternative strategies

Transcriptome assemblers are reported to be capable of outputting complete or nearly complete transcripts for different splicing isoforms (Grabherr *et al.*, 2011). If the transcripts were complete, Glutton would not improve upon even simple heuristics such as selecting the longest contig. Many transcripts are lowly expressed, however, and we expect Glutton to show substantial improvements over heuristics. We compared the output of Glutton with two commonly used strategies for post-processing transcriptome data prior to downstream analysis: selecting the longest contig (Longest contig) and selecting the contig with the highest identity to a reference gene (Highest identity). We processed the same four *D. melanogaster* samples used previously together with all reference species from Ensembl Metazoa. Data was generated by extracting individual contigs from the PAGAN alignments produced by Glutton’s align command. These contigs were post-processed in the same manner as before: trimming at stop codons and rejecting alignments based on insufficient coverage (90%), identity (30%) and overlap (100 bp).

The results for sample SRR166808 using *D. grimshawi* as a reference species are shown in Table 2. For all assemblers, Glutton outperforms both heuristics in terms of total alignments, in the case of Trinity, for example, Glutton outputs 20% more alignments than Longest contig. Importantly, the higher number of alignments does not significantly affect Glutton’s accuracy, which remains comparable to both heuristics, the maximum difference being 2.2% (to Highest identity using Oases). The results are similar with the other three RNA-seq data sets and, for all samples tested, Glutton outputs more alignments than both heuristics irrespective of transcriptome assembler or reference species (median 21% more alignments, IQR: 12-33%, data not shown). Although Highest identity outperforms others in terms of accuracy, Glutton’s accuracy is comparable with a median difference across all experiments of 1.8%.

**Table 2:** Transcriptome assembler and Glutton statistics for sample SRR166808 with *D. grimshawi* as reference species. As well as Glutton’s depth-based approach we compared the effect of taking the longest contig and the contig with the highest identity to a reference gene. *\** = rerun as single-ended reads.

|  | Glutton |  | Longest contig |  | Highest identity |  |
| --- | --- | --- | --- | --- | --- | --- |
|  | Alignments | Correct (%) | Align. | Corr. | Align. | Corr. |
| TransABySS | **6,718** | 92.1 | 5,879 | 93.0 | 4,982 | **93.6** |
| Oases | **6,604** | 88.7 | 4,739 | 90.1 | 3,893 | **90.9** |
| Trinity | **6,721** | 92.7 | 5,549 | 93.4 | 5,446 | **93.6** |
| SOAPdenovo-Trans | **5,979** | 94.2 | 5,430 | 94.8 | 5,497 | **94.9** |
| SOAPdenovo-Trans* | **6,878** | 93.3 | 5,512 | 94.5 | 5,367 | **94.8** |

### 2.4 Comparison with reference transcripts

Glutton is designed to process transcriptome assemblies for use with downstream evolutionary analysis methods. RNA-seq targets coding regions in the genome and one relevant analysis for the assembled transcripts is detection of positive Darwinian selection affecting protein-coding genes. To understand how Glutton’s RNA-seq data-derived alignments perform in such analyses, we contrasted estimates of synonymous and non-synonymous substitution rates for codon sequences between Glutton alignments and reference alignments from EnsemblCompara.

We retrieved RNA-seq data from seven *Drosophila* species: *D. ananassae, D. melanogaster, D. mojavensis, D. pseudoobscura, D. simulans, D. virilis* and *D. yakuba* (accessions SRR768437, SRR166809, SRR166834, SRR166830, SRR166813, SRR768439 and SRR768435, respectively). Each sample was filtered for quality and assembled with SOAPdenovo-Trans using single-end reads. Raw reads were mapped to assemblies using BWA. All seven samples were processed with Glutton using *D. grimshawi*, *Z. nevadensis* and M.*leidyi* as reference species from Ensembl Metazoa (release 25). After splitting each gene family alignment on paralogs from the reference species, Glutton output 8,119 alignments with *D. grimshawi* as reference species, 3,369 with *Z. nevadensis* and 1,364 with *M. leidyi*. Figure 6 breaks down the total number of alignments by the number of species present in each. Using *D. grimshawi* as the reference, for example, output 1,757 alignments containing at least six *Drosophila* species.

**Figure 6.**
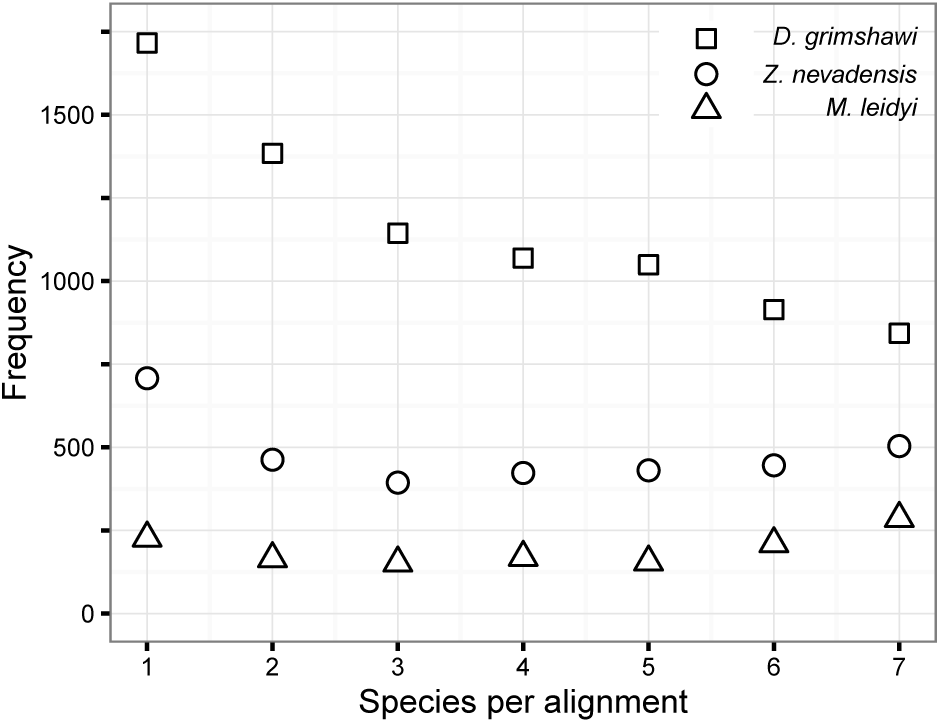
Breakdown of the number of species represented in Glutton alignments of seven *Drosophila* species.

Our goal was to compare *dN /dS* ratios produced by an alignment derived from RNA-seq data with that given by the complete transcript data from EnsemblCompara. We removed the reference genes, discarded any alignments containing less than six species and, per sub-alignment, inferred a phylogenetic tree using RAxML (Stamatakis, 2014). For comparison, alignments and phylogenetic trees from those species present in a given Glutton alignment were retrieved from EnsemblCompara. Both Glutton and Ensembl alignments were analysed using PAML (Yang, 2007) and SLR (Massingham and Goldman, 2005) to calculate *dN /dS*. We noticed that alignments from Ensembl produced higher than expected *dN /dS* ratios (see Supplemental data, Figure S7), so reference sequences were realigned with PAGAN using a translated alignment with the EnsemblCompara phylogeny as guide tree.

Glutton shows high concordance with results based on complete reference transcripts from EnsemblCompara irrespective of divergence time from the reference species. Figure 7 shows *dN /dS* from PAML assuming the same ratio over all sites (model M0). The alignments derived from using *D. grimshawi* as a reference are of high quality and show a similar spread compared to the alignments from EnsemblCompara. As divergence times increase, however, those genes that are output tend to be more conserved as evidenced by identifying only genes undergoing stronger purifying selection. While this exemplifies the need for a suitable, closely related reference species, even the results using more distantly related reference species, though limited, can still be trusted if nothing else is available. The results for SLR are presented in Supplemental materials, Figures S8 and S9.

**Figure 7.**
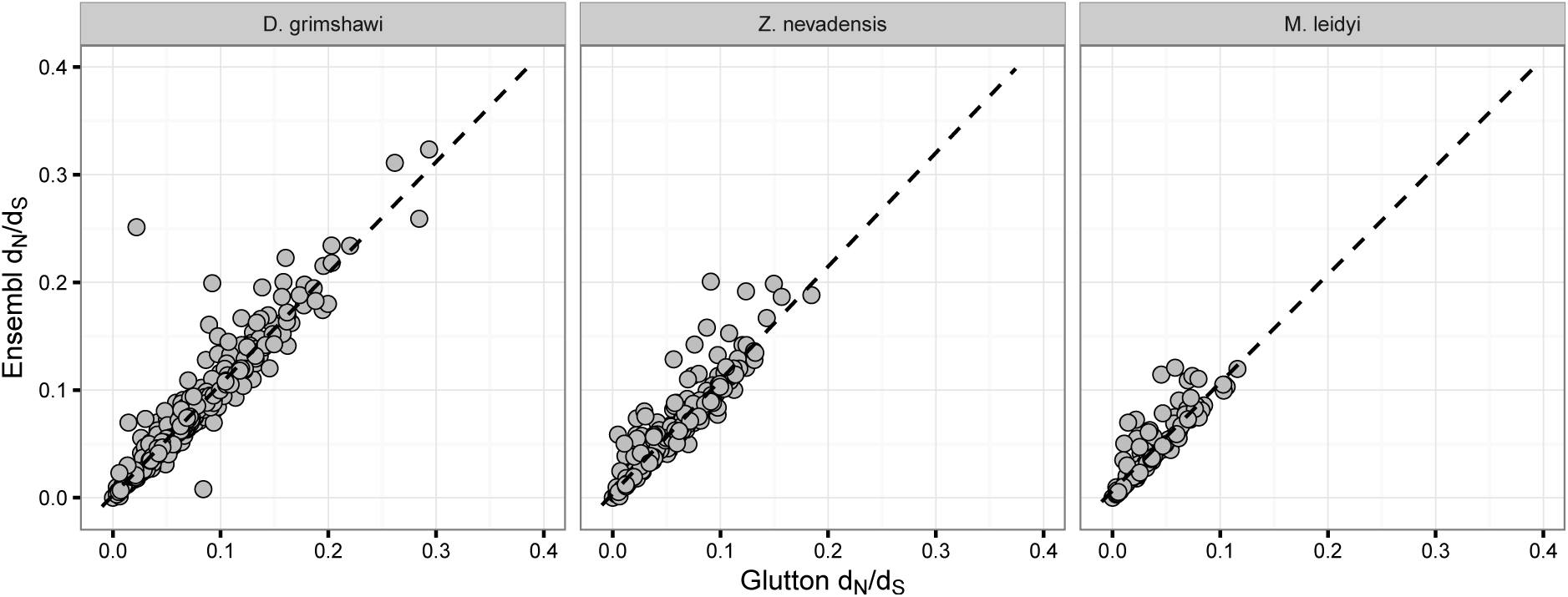
Comparison of *dN /dS* values (PAML, model M0) for seven *Drosophila* species using three different reference species. Dashed line shows line of best fit.

### 2.5 Application to thermal adaptation in Malagasy dung beetles

We applied Glutton to the study of thermal adaptation in large-bodied dung beetles from Madagascar, whose lineage is composed of twelve abundant and ecologically successful species. Species in this lineage are morphologically and ecologically similar differing mainly in their elevation distributions, with six species (*N. vadoni*, *N. manomboensis*, *N. binotatus*, *N. bimaculatus*, *N. antsihakanensis* and *N. humbloti* “vadoni” clade) occurring at low elevations (0–500m) and the other six (*N. viettei*, *N. pseudoviettei*, *N. dubitatus*, N.*nitens*, *N. clypeatus* and *N. mirjae* “viettei” clade) restricted to mid/high elevations (500–1500m) (Montreuil *et al.*, 2014; Miraldo *et al.*, 2011; Miraldo and Hanski, 2014). Recent research involving three of these species (*N. vadoni*, *N. dubitatus* and *N. nitens*) showed significant differences in species’ thermal tolerances, with the low-elevation species (*N. vadoni*) showing the highest critical thermal maximum (CTmax) and the high-elevation species (*N. nitens*) the lowest, and significant variation also detected among populations within species (Va¨litalo, 2013). These results suggest that species in this group might be thermally adapted to the different climatic conditions along elevational gradients, and are therefore an ideal model to test the role of thermal adaptation in diversification and local adaptation.

Nine *Nanos* species composed of 22 samples were analyzed using pooled RNA-seq data (the 22 samples were composed of 287 individual beetles, see Table S1 in Supplemental materials for details). The samples were collected and sequenced using standard procedures (see Supplemental Methods). In addition to the *Nanos* species we included bull-headed dung beetle (also known as taurus scarab) *Onthophagus taurus* (Misof *et al.*, 2014) for use as an outgroup. Each sample was filtered for quality, assembled with SOAPdenovo-Trans using single-end reads and the raw reads mapped back to the assemblies using BWA. All 23 samples were processed with Glutton using *Tribolium castaneum* as reference species from Ensembl Metazoa (release 25) which, when split on paralogs from the reference species, produced 6,162 multiple sequence alignments that contained at least one beetle species. We removed the reference genes, discarded any alignments containing less than 7 species and, per sub-alignment, performed a realignment using PRANK and inferred a phylogenetic tree using RAxML. The resulting 3,038 subalignments and phylogenetic trees were analysed using SLR, which resulted in 51 genes identified as under putative selection after correcting for multiple comparisons. This list of genes was compared to candidate genes thought to be involved in thermal adaptation in *Drosophilia* (Hangartner *et al.*, 2015), but no known candidates were identified.

An analysis based on an earlier assembly with Trinity had suggested that *Hypoxia up-regulated protein* gene (TC007793 in *T. castaneum*) was under strong positive selection. However, the region includes large indels in two *Nanos* species and, when the original data were mapped to the scaffolds, the sequencing coverage significantly dropped around the indels (Figure 8a). The analysis with SOAPdenovo-Trans, however, showed that this cluster of sites under positive selection was an artefact of incorrect assembly (Figure 8b). This error was relatively easy to detect due to the high coverage, but it would be difficult to correct in a consistent manner without risking excessive false positives in lowly expressed transcripts.

**Figure 8:**
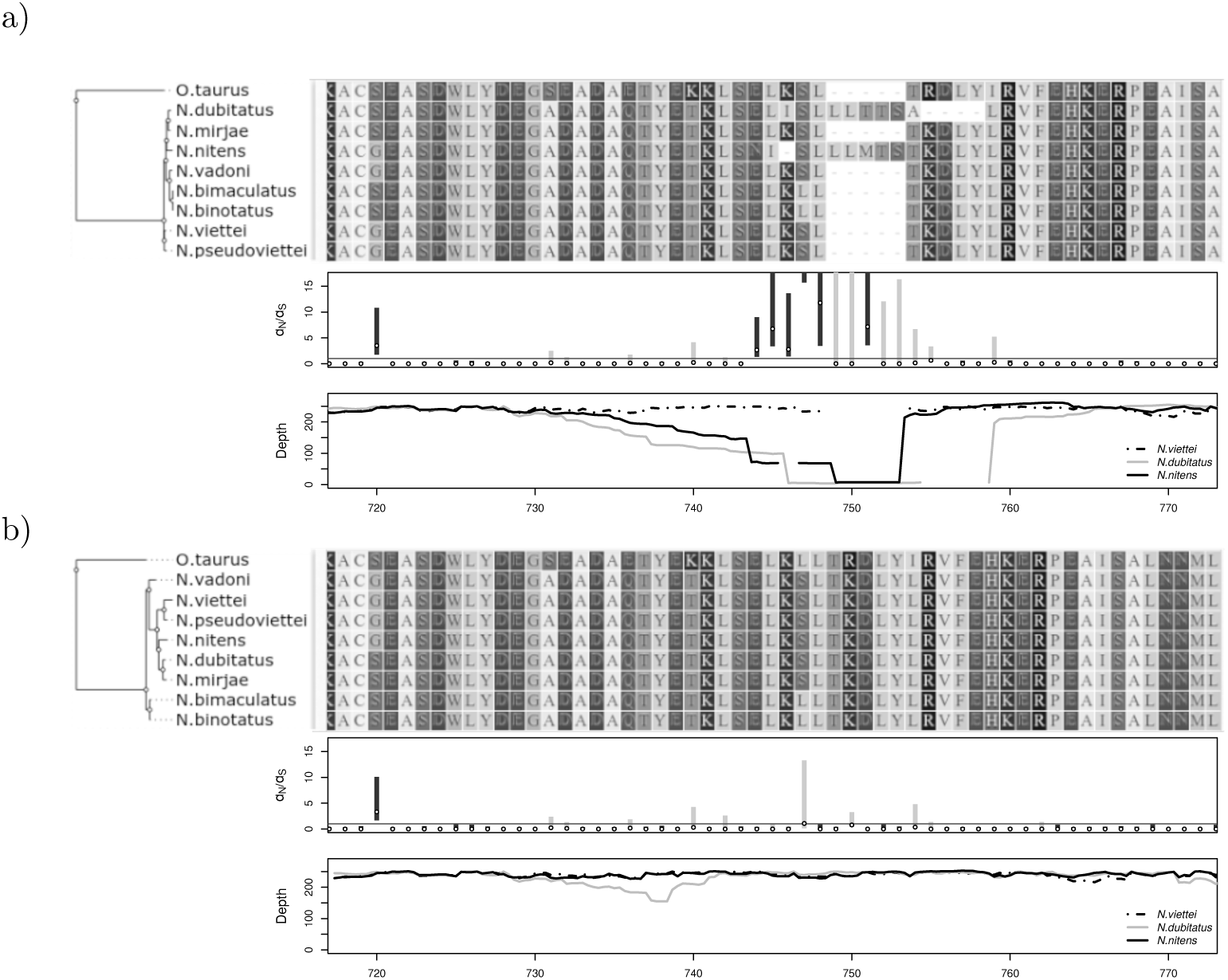
Selection analysis of dung beetle *Hypoxia up-regulated protein* gene using (a) Trinity assembly and (b) SOAPdenovo-Trans assembly. (a) The alignment (top panel) shows overlapping indels around position 750. The selection analysis with SLR (middle panel) identifies several apparently positively selected sites in this region, indicated by *d*n*/d*s point estimates (circles) and error bars (gray bars) above 1. The local dips in read depth (bottom panel) suggest that the assemblies containing the indels are erroneous and do not represent the dominant allele. (b) A reanalysis with a different assembly shows no indels or positively selected sites in the region. Alignment and tree were visualized with Wasabi (Veidenberg *et al.*, 2015) and other graphics generated using R (R Core Team, 2013) and Rsamtools (Morgan *et al.*, 2015).

## 3 Discussion

Glutton produces high quality gene alignments across a comprehensive range of divergence times between study and reference species. The selection of an appropriate reference species is critical and more closely related species, as expected, perform substantially better than distant ones in terms of the number of alignments output and the proportion of those alignments that are correct in terms of identifying correct orthologs. The choice should not always be made by minimising the divergence time, however, as some reference species systematically produce more erroneous alignments than predicted, for example, we identified *M. cinxia* as a poor reference for use with Glutton (Figure 5). Its weak performance is not explained by evolutionary rate or gene content only as our reference set contains three other Lepidopteran species with a shared history and similar gene numbers (16,639 in *M. cinxia* vs. 12,669-16,254 in other Lepidopteran species). While we cannot predict the performance of a specific reference (the number of error-free alignments being unknown), it is clear that quality varies significantly and reported genes can be incomplete, erroneous or have incorrect gene family memberships. Taking this into account, if the most closely related reference is less well studied, we advise users to supplement their analysis by separately using a more distant, better studied reference and comparing results.

Glutton’s performance is not only determined by the reference species, the study species is also a factor, specifically related to gene count. In our analyses, Glutton performed better, the fewer genes in the study species. Of the species we studied, the one with the highest number of genes, *A. pisum*, had the highest proportion of erroneous results. It is an exceptional species, however, having both lost genes essential to most other organisms and undergone substantial gene family duplication. While Glutton can differentiate between which paralog is most suitable from the reference species, in the case of many-to-one relationships, where several transcripts are orthologous to the same reference gene, they will be erroneously merged. One of our future goals is to detect when this happens and distinguish between paralogs in the study species assigned to the same reference gene without being misled by assembly errors.

We have shown that Glutton works well with all the transcriptome assemblers tested. Most transcriptome assemblers infer alternative splicing isoforms along with the assembly, which we had assumed would provide additional context compared to using contigs alone, producing better BLAST hits and, therefore, superior results. However, two of the best performing methods (SOAPdenovo-Trans with single reads and TransABySS) only produce contigs and no isoforms. Indeed, depending on the method by which isoforms were derived, they appear detrimental: the two worst performing methods (Oases and SOAPdenovo-Trans with paired-end data) use the same method to enumerate all transcripts. We believe that while the correct transcripts are present, the inclusion of so many other possible versions of the same gene make the alignment extension problem more complex and degrades performance.

Glutton produces higher numbers of alignments than common heuristics for selecting the best contigs for downstream analysis, without sacrificing accuracy. For comparative analysis, multiple sequence alignments output by Glutton give results that are concordant with alignments using complete reference data. While the accuracy of downstream analyses based on Glutton-generated alignments did not appear to be affected by divergence time, the total yield is reduced with distant references as only conserved genes can be reliably extracted from assemblies. This is hardly surprising, but even the *∼*500 gene alignments containing at least six of the seven *Drosophila* species using the most distant reference in our analysis is beyond what could have been imagined only a few years ago. Indeed, our analyses on fruit flies and dung beetles confirm that the set of highly expressed genes in whole body samples are very similar between closely related species (Hittinger *et al.*, 2010).

Glutton is similar in operation to Scaffolding using Translation Mapping (STM), which also uses reference data to orientate and merge transcripts homologous to the same gene (Surget-Groba and Montoya-Burgos, 2010). STM predates many of the transcriptome assembly programs tested, with most supporting the multi-*k*-mer approach it introduced. Unlike Glutton, STM is focused on producing an optimised assembly and does not output multiple sequence alignments. Moreover, STM does not take sequencing depth into consideration, limiting it to arbitrary tie breaks in the case of contigs mapping to the same loci. In a similar vein, several *de novo* assemblers are capable of performing guided assembly; mapping reads to a closely related reference species to bootstrap assembly (e.g. the Columbus module for Velvet (Zerbino, 2010) and Trinity support guided assembly). These methods were not included in our analysis as they only work with very close reference species. Another recent approach called Direct Genome Mapping (DGM) also advocates mapping RNA-seq data directly to a closely related species, purporting to show an improvement over *de novo* assembly for use in functional annotation (Ockendon *et al.*, 2015). DGM is only demonstrated with very closely related species (diptera and primates) which, for many study species, is impractical. While their results appear impressive, unlike Glutton, their assembly-based methods do not take advantage of translation and require an exceptionally conservative E value of 1 *×* 10*−*10 to be considered homologous: a level of significance their own method need not pass. We found that Glutton can retrieve a significant proportion of *D. melanogaster* genes over a comprehensive range of divergence times (see Supplemental materials, Figures S5, S6) and consider their results to be overly pessimistic for *de novo* assembly-based approaches.

Future developments that we intend to investigate include the use of multiple reference species and the incorporation of complete gene models. Using multiple reference species would enable Glutton to output more alignments due to increased sensitivity from having a greater range of reference sequences. Problems may arise, however, in the case of genes that are highly conserved across many species; transcripts from two species with superficial differences may be most similar to different genes resulting in unwanted fragmentation. This issue might be solved by preferring inferred ancestral sequences over extent sequences. With respect to gene models, we currently use canonical transcripts to build a reference. To use gene models, PAGAN would need to be extended to allow for extant tree nodes to be graphs encoding known gene models and Glutton changed to extract this information from Ensembl. It is possible that incorporating gene models, where available, would improve sensitivity as we are currently limited by what transcripts are highly expressed in a given sample. Information on alternative splicing in non-model species, such as *T. castaneum* in our dung beetle analysis, is limited and the quality of gene models can be poor. Furthermore, alternatively spliced exons may not be as well conserved across distant species as constitutive ones, and they are not necessarily expressed in significant quantities in bulk tissue under normal conditions. It is therefore unclear if in aggregate this would give a significant improvement over our current approach.

## 4 Materials and Methods

Glutton is composed of three subcommands: *build* (responsible for creating reference databases using EnsemblCompara), *align* (assigning contigs to reference transcripts and reference alignment extension) and *scaffold* (merging alignments into scaffolds and consensus alignments).

### 4.1 Constructing the reference database

Glutton requires a database of reference transcripts together with orthology information. Data is retrieved using the programmatic interface to the BioMart service at Ensembl to download all CDS sequences for a given species (Kinsella *et al.*, 2011). Homology information is retrieved using the Ensembl REST API (Yates *et al.*, 2014). Gene families containing two or more genes are aligned with PRANK using the translated alignment option. The resulting multiple sequence alignment, together with the phylogenetic tree estimated by neighbour-joining are packaged into a single compressed database file.

### 4.2 Gene family assignment and alignment extension

All reference transcripts from the Glutton database are used to create a BLAST protein database. Contigs are assigned to reference transcripts using BLASTX. Queries are split into batches and executed in parallel. BLAST hits are filtered for identity, length and E value (default thresholds: 0.3, 100 bp, 1 *×* 10*−*3, respec-tively). Contigs are assigned to the gene family containing the top BLAST hit. Glutton takes each gene family for which contigs were assigned and uses PAGAN extend the reference alignment from the database using assigned contigs as query sequences. All alignments are executed in parallel. If multiple assemblies from multiple species are being processed simultaneously, users must provide a *sample* and a *species* ID for each assembly.

### 4.3 Scaffolding and post-processing

The scaffolding command takes alignments produced by PAGAN and merges them using the metadata provided by the user. One BAM file can be provided per assembly for sequencing depth information. To perform scaffolding, contigs with the same sample ID are merged if they are aligned to the same reference transcript and, if they overlap, there is only mismatches between the two contigs. Contigs identified by the assembler as isoforms of the same gene are not merged, however, this feature is only available for Trinity, Oases and SOAPdenovo-Trans. To construct each consensus alignment, contigs with the same species ID are ordered by average sequence depth and merged sequentially from lowest to highest, allowing contigs with greater depth to overwrite those with lower. Output consensus alignments are trimmed at the 5’ end at the start codon (or at the same column as the reference transcript start codon, if missing) and at the 3’ end at the first stop codon. Consensus alignments are filtered for protein identity and overlap with the reference transcript (defaults: identity 0.3, overlap 0.9).

### 4.4 Data access

RNA-seq samples from Malagasy dung beetle species can be accessed under BioProject PRJNA314439.

## 5 Acknowledgements

This work was supported by the Marie Curie Career Integration Grant to A.L.. We acknowledge CSC - IT Center for Science, Finland, for computational resources, and Baylor College of Medicine Human Genome Sequencing Center for the *Onthophagus taurus* RNA-seq data.

